# A joint metabolic–integrative regime of consciousness: evidence from FDG-PET and TMS–EEG

**DOI:** 10.64898/2026.04.18.719419

**Authors:** Thyagarajan Shivashanmugam, Anish Mehta

**Author notes:** **Corresponding Author** Thyagarajan Shivashanmugam, Chief Scientific Officer, Placeboes Research Foundation, Bengaluru, India.

## Abstract

Conscious experience is associated with both sufficient cerebral metabolic support and preserved large-scale neural integration, yet these two constraints have largely been studied in isolation. Whether consciousness requires their *joint* satisfaction — and whether either alone is sufficient — remains an open question. Here, we test the prediction that conscious states occupy a joint metabolic–integrative regime defined by the co-occurrence of adequate metabolic support and preserved perturbational complexity.

We performed a constraint-driven cross-study synthesis of published benchmarks from [^18^F]- fluorodeoxyglucose positron emission tomography (FDG-PET) and transcranial magnetic stimulation combined with electroencephalography (TMS–EEG). Metabolic values (expressed as percentage of normal cortical metabolism) and perturbational complexity index (PCI) values were mapped across canonical states spanning disorders of consciousness, sleep, and anaesthesia into a shared two-dimensional state space.

A metabolic lower bound of ∼42–46% of normal cortical metabolism and a complexity threshold of PCI* ≈ 0.31 consistently separated unconscious from conscious conditions across independent cohorts. Of 15 conditions with complete data on both axes, all 9 conscious states occupied the joint regime (above both thresholds) and all 6 unconscious states fell below at least one threshold (Fisher’s exact test, two-tailed p ≈ 2.00 × 10^−4^); metabolic support and perturbational complexity were strongly correlated across conditions (Spearman’s ρ = 0.95). The sole off-diagonal condition — NREM sleep, with preserved metabolism (∼56% of normal) but low complexity (PCI ≈ 0.23) — was unconscious, consistent with the prediction that metabolic support is necessary but not sufficient without preserved integrative dynamics.

These findings support a joint metabolic–integrative constraint on conscious experience. The framework yields a clear falsification target: reportable experience should not occur in conditions falling outside this joint regime. Prospective within-subject studies combining FDG-PET and TMS–EEG are needed to evaluate this prediction at the individual level.

## Introduction

Identifying the biological conditions that support conscious experience remains a central challenge in neuroscience (Seth and Bayne, 2022). While substantial progress has been made in characterising neural correlates of consciousness (Tononi, 2004; Koch *et al*., 2016), many approaches have focused on single dimensions of brain function. In particular, measures of large-scale integration and information complexity have demonstrated strong discriminative power between conscious and unconscious states (Casali *et al*., 2013; Casarotto *et al*., 2016; Comolatti *et al*., 2019), whereas metabolic studies have identified thresholds associated with recovery of awareness (Stender *et al*., 2014, 2016). However, these lines of work have largely developed in parallel, and it remains unclear whether either dimension alone is sufficient or whether conscious states depend on the joint satisfaction of multiple biological constraints.

We have proposed a biologically grounded framework (Shivashanmugam and Mehta, 2026) in which conscious experience arises under constraints imposed by both metabolic support and large-scale neural integration. In this formulation, conscious states are predicted to occupy a joint regime defined by sufficient energetic availability and preserved perturbational complexity. A key implication of this framework is that dissociations between these dimensions—configurations in which metabolic support and integrative dynamics are not aligned—should be rare, and the presence of verified experience under conditions of reduced metabolism or reduced perturbational complexity would constitute a falsifying observation.

Two empirical domains provide complementary access to these constraints. Measures of cerebral glucose metabolism derived from [^18^F]-fluorodeoxyglucose positron emission tomography (FDG-PET) index the energetic support available to cortical processes and distinguish vegetative state/unresponsive wakefulness syndrome (VS/UWS) from minimally conscious state (MCS) (Stender *et al*., 2016; Hermann *et al*., 2021), supporting the existence of a metabolic boundary associated with awareness. In parallel, perturbational complexity measures derived from transcranial magnetic stimulation combined with electroencephalography (TMS–EEG), particularly the perturbational complexity index (PCI) (Casali *et al*., 2013), quantify the capacity of the cortex to sustain integrated yet differentiated responses and reliably separate conscious from unconscious conditions across wakefulness, sleep, anaesthesia, and disorders of consciousness (Massimini *et al*., 2005; Casarotto *et al*., 2016; Comolatti *et al*., 2019).

Despite the robustness of these measures, their relationship has not been systematically evaluated within a common framework. In particular, it remains unresolved whether metabolic support and perturbational complexity represent independent correlates or whether conscious states are constrained to a region in which both conditions are satisfied.

Here, we evaluate this prediction by synthesising cross-study benchmarks from FDG-PET and TMS–EEG within a two-dimensional state space defined by metabolic support and perturbational complexity. In doing so, we (i) operationalise empirically anchored boundaries on each axis, (ii) assess whether canonical brain states cluster within a jointly sufficient regime, and (iii) delineate falsification-relevant “off-diagonal” configurations for future within-subject testing.

## Methods

### Study design and analytical framework

We conducted a constraint-driven cross-study synthesis (non-meta-analytic) to evaluate a predefined theoretical prediction: that reportable conscious states occupy a regime defined by the joint presence of sufficient metabolic support and preserved perturbational complexity. This prediction was operationalised within a two-dimensional state space comprising (i) a metabolic axis, indexed by cortical glucose metabolism derived from FDG-PET, and (ii) a complexity axis, indexed by perturbational complexity measured using TMS–EEG, primarily the perturbational complexity index (PCI).

States were treated as observational units and indexed either by clinical diagnosis such as vegetative state/unresponsive wakefulness syndrome [VS/UWS], minimally conscious state [MCS], locked-in syndrome [LIS] (Giacino *et al*., 2002) or by experimentally induced condition (e.g., propofol-induced unresponsiveness, sleep stages, ketamine exposure). The analytical objective was to evaluate cross-study convergence within the defined state space rather than to perform quantitative meta-analytic pooling.

Two classes of configurations were defined a priori as falsification-relevant: (i) preserved metabolic support with reduced perturbational complexity, and (ii) preserved perturbational complexity under insufficient metabolic support.

### Data sources and eligibility

The synthesis was restricted to peer-reviewed cohort studies reporting quantitative measures of cerebral glucose metabolism (FDG-PET) and/or perturbational complexity (TMS– EEG/PCI) in conditions relevant to disorders of consciousness, sleep, and anaesthesia.

Eligible studies comprised two categories: (i) within-cohort multimodal studies reporting paired metabolic and electrophysiological measures, and (ii) single-modality benchmark studies used to anchor each axis. Inclusion required reporting of quantitative values or thresholds sufficient to position observations within the defined metabolic–complexity state space.

Conditions for which quantitative values were available on only one axis were retained in descriptive tables but excluded from the joint-quadrant classification and statistical analyses. Specifically, midazolam deep sedation and xenon anaesthesia were excluded from the joint analysis due to the absence of quantitative FDG-PET metabolic values in the available literature; a further sedation condition (Laaksonen *et al*., 2018) was excluded due to the absence of PCI data.

### Study identification

Relevant studies were identified through targeted searches of PubMed and reference lists of key articles using combinations of terms including “FDG-PET”, “metabolism”, “TMS–EEG”, “PCI”, “disorders of consciousness”, “sleep”, and “anaesthesia”. Searches were focused on identifying benchmark studies reporting quantitative values sufficient to anchor the metabolic and complexity axes.

Study selection was guided by relevance to the predefined analytical framework and by the availability of quantitative measures enabling placement within the metabolic–complexity state space. Priority was given to widely cited datasets establishing empirical thresholds or distributions across conditions.

### Threshold definition and interpretive approach

Two empirically derived boundaries were used to anchor the joint-regime analysis. On the metabolic axis, prior FDG-PET studies identify a lower bound near ∼42% of normal cortical metabolism in VS/UWS (Stender *et al*., 2016), with higher values (∼55–60%) observed in MCS (Stender *et al*., 2016). On the complexity axis, the principal operational threshold was PCI* ≈ 0.31, derived using receiver operating characteristic analysis of PCImax values across benchmark conscious and unconscious states (Casarotto *et al*., 2016). These thresholds were not re-estimated. Instead, they were treated as externally derived reference points, and the analysis evaluated cross-study consistency with the joint-regime prediction. For the quadrant-based classification, a single operational threshold of ∼44% of normal cortical metabolism was applied. This value represents the lower bound of the transition band (∼42–46%) identified across independent FDG-PET datasets (Stender *et al*., 2016) and is used here as a conservative cutoff rather than a precise biological boundary. The band rather than a point estimate reflects the graded nature of the VS/UWS–MCS boundary in the source data.

### Metabolic axis

Metabolic support was indexed using FDG-PET measures of cerebral glucose consumption. Where available, normalised cortical metabolism expressed as a percentage of healthy-control values was used directly. For studies reporting the metabolic index of the best-preserved hemisphere (MIBH), published mappings to percentage-of-control metabolism were applied to ensure comparability (Hermann *et al*., 2021).

In pharmacological conditions, absolute or relative changes in whole-brain glucose metabolic rate were extracted where reported. These measures were interpreted as directional indicators of energetic modulation and were not assumed to be directly comparable to normalised percentage-of-control values unless explicitly calibrated within the source study.

To enable cross-study integration, metabolic measures were interpreted on a common conceptual scale anchored to percentage-of-normal cortical metabolism. Empirical reference points derived from disorders of consciousness — particularly the separation between VS/UWS (∼42%) and minimally conscious state (∼55–60%) — were used to position observations along the metabolic axis (Table 1) (Stender *et al*., 2016).

**Table 1.** FDG-PET Metabolic Values by Consciousness State.

| Table 1. FDG-PET Metabolic Values by Consciousness State |  |  |  |  |  |  |
| --- | --- | --- | --- | --- | --- | --- |
| State | Metabolism (% of normal) | Source | Absolute CMRGlu | MIBH | Sample (n) | Conscious classification |
| Healthy wakefulness (reference) | ~100% (by definition; normative baseline) | Stender et al., 2016 | $29 \pm 8 \mu\text{mol} \cdot 100\text{g}^{-1} \cdot \text{min}^{-1}$ (awake GMR) | — | n=6 (Alkire 1995); n=29 controls (Stender 2016) | Conscious |
| MCS | 55% global cortical CMRglc; ~60% frontoparietal | Stender et al., 2016 | — | MIBH $4.16 \pm 1.11$ | n=21 MCS (Stender); n=87 MCS (Thibaut 2021) | Conscious |
| EMCS | ~55% (indistinguishable from MCS in source) | Stender et al., 2016 | — | — | n=6 EMCS | Conscious |
| LIS | Near-normal (gray-matter SUV $5.6 \pm 3.2$; not expressed as % of normal)† | Soddu et al., 2015 | — | — | n=4 LIS | Conscious |
| REM sleep | ~100% of waking (global metabolic rate not significantly different from waking: $+4.30 \pm 7.40\%$, $p = 0.35$)‡ | Maquet et al., 1990 | — | — | n=9 | Conscious |
| Ketamine (subanesthetic) | Increased $+11.9 \pm 10.9\%$ vs baseline (above normal) | Långsjö et al., 2004 | — | — | n=9 | Conscious (dissociative) |
| VS/UWS | 42% global cortical CMRglc | Stender et al., 2016 | — | — | n=14 (Stender); n=48 (Thibaut 2021) | Unconscious |
| VS/UWS (Liu cohort) | 41% (metabolic index $3.32 = 41\%$ of healthy mean $7.88$) | Liu et al., 2024§ | — | — | n=48 VS/UWS | Unconscious |
| NREM sleep | ~56% of waking (global rate reduced by $-43.80 \pm 14.10\%$, $p < 0.01$)‡ | Maquet et al., 1990 | — | — | n=9 | Unconscious |
| Propofol anaesthesia | ~45% of normal (awake GMR $29 \pm 8$; anaesthetised $13 \pm 4 \mu\text{mol} \cdot 100\text{g}^{-1} \cdot \text{min}^{-1}$; ratio = 45%)* | Alkire et al., 1995 | $13 \pm 4 \mu\text{mol} \cdot 100\text{g}^{-1} \cdot \text{min}^{-1}$ | — | n=6 | Unconscious |
| Midazolam deep sedation | No FDG-PET metabolic value available in literature** | — | — | — | n=6 (PCI only; Casali 2013) | Unconscious |
| Xenon anaesthesia | No FDG-PET metabolic value available in literature** | — | — | — | n=6 (PCI only; Casali 2013) | Unconscious |
| Sevoflurane anaesthesia | ~61% of normal (~39% reduction; not expressed as % of normal in source) | Jeong et al., 2006 | — | — | n=8 | Unconscious |
| † LIS metabolic value not expressed as % of normal in source; near-normal metabolism assumed for state-space placement based on preserved cortical integrity in LIS.<br>‡ REM and NREM metabolic values from Maquet et al. (1990), which used [<sup>18</sup>F]FDG PET; values represent global cortical metabolic rate relative to waking baseline in the same subjects. § Liu et al. published 2024 (DOI: 10.3389/fneur.2024.1425271); some databases index publication year as 2025. *** Propofol % of normal derived from absolute CMRGlu ratio (13/29 = 45%); not directly normalised to a separate healthy-control group in the source study. ** Midazolam and xenon excluded from joint-quadrant classification due to absence of FDG-PET metabolic data; PCI values available in Table 2. |  |  |  |  |  |  |

### Complexity axis

Perturbational complexity was indexed using PCI values derived from TMS–EEG studies. The operational boundary separating conscious from unconscious conditions was defined as PCI* ≈ 0.31, corresponding to the empirically derived threshold distinguishing states with and without subjective report (Casarotto *et al*., 2016).

PCI values were extracted across canonical physiological and clinical states, including wakefulness, sleep, anaesthesia, and disorders of consciousness (Casali *et al*., 2013; Casarotto *et al*., 2016). Particular emphasis was placed on (i) VS/UWS subgroups exhibiting PCI values above the threshold (Casarotto *et al*., 2016) and (ii) clinically conscious but behaviourally constrained conditions, such as locked-in syndrome, in which preserved complexity is expected (Casali *et al*., 2013).

Each condition was represented using reported metabolic and perturbational complexity values positioned relative to externally derived thresholds, enabling direct comparison within a common two-dimensional (M, C) state space (Figure 1). For the joint state-space representation, each condition is represented as a coordinate pair (M, C), where M denotes normalised metabolic support (percentage of normal cortical metabolism) and C denotes perturbational complexity (PCI value relative to PCI* threshold).

**Figure 1.**
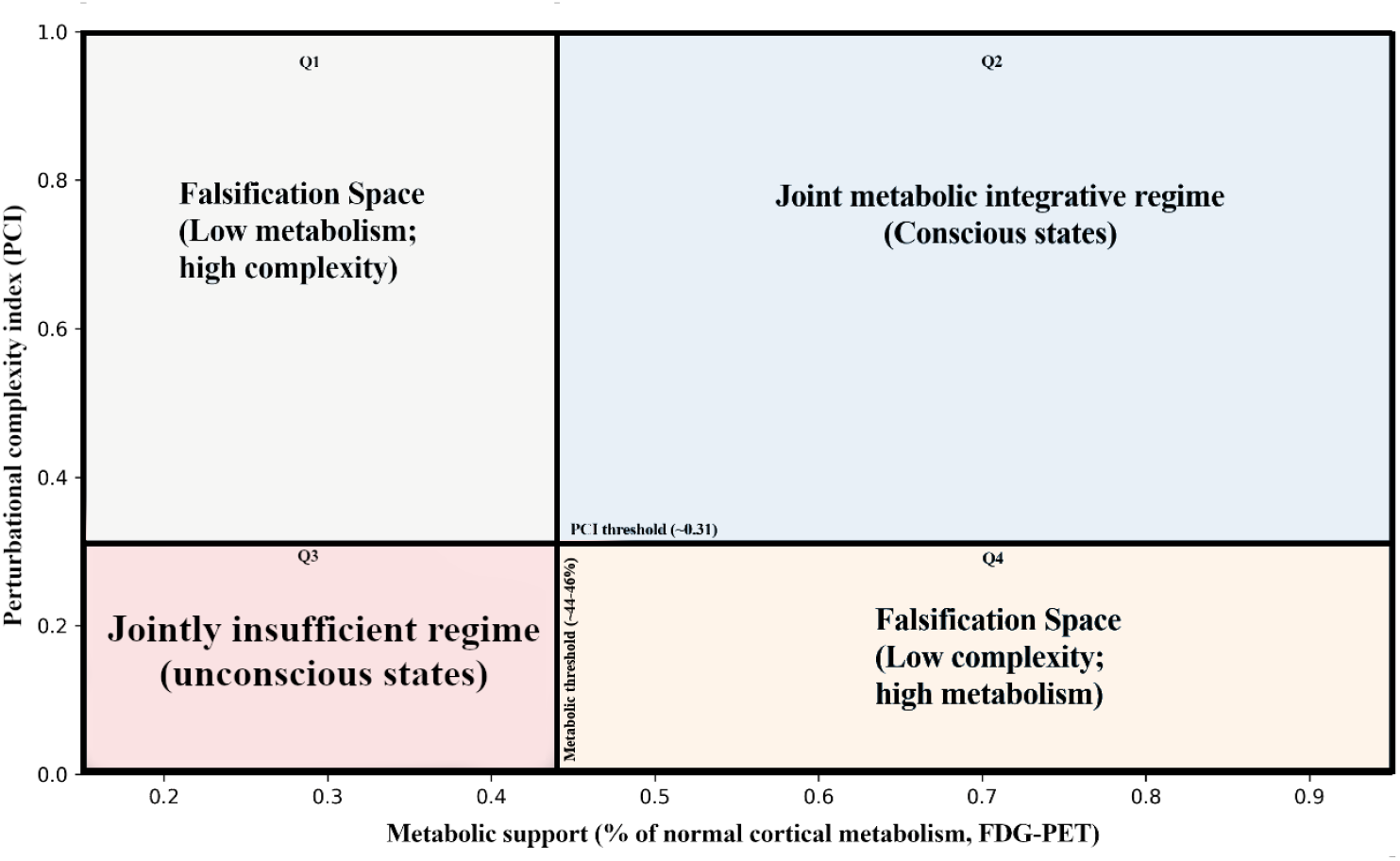
Joint metabolic–integrative state space. Joint state space defined by metabolic support (M; % of normal cortical metabolism, FDG-PET) and perturbational complexity (C; PCI). Thresholds at ∼44–46% metabolism and PCI* ≈ 0.31 delineate four quadrants: Q2, the joint metabolic–integrative regime (conscious-compatible); Q3, a jointly insufficient regime (unconscious); and Q1 and Q4, off-diagonal configurations representing falsification-relevant regions.

### Cross-study harmonisation

FDG-PET–derived metabolic support was harmonised by expressing all values as percentage of normal cortical metabolism relative to healthy age-matched controls. For studies reporting the metabolic index of best-preserved hemisphere (MIBH), published mappings to percentage-of-control metabolism were applied (Hermann *et al*., 2021). When anaesthesia studies reported absolute CMRGlu or relative changes rather than normalised percentage-of-control values, these measures were treated as directional indicators on the metabolic axis and positioned relative to disorders-of-consciousness benchmarks (Stender *et al*., 2016) rather than assumed to be directly numerically comparable. This reliance on external calibration points introduces cross-study variability that should be considered when interpreting cross-state placements.

Where source studies reported metabolic ranges rather than point estimates (e.g., VS/UWS ∼38–42%), the midpoint of the reported range was used as the representative coordinate for state-space placement. For conditions reported across multiple independent cohorts, each cohort-level estimate was retained as a separate entry in the state-space mapping to preserve cross-study variability rather than averaging across datasets.

TMS–EEG–derived perturbational complexity was harmonised by retaining only PCI values computed using published algorithms: the original PCI (Casali *et al*., 2013) or the state-transition variant PCIst (Comolatti *et al*., 2019). The operational threshold PCI* ≈ 0.31 was derived via receiver operating characteristic analysis across a large benchmark dataset encompassing multiple stimulation sites and recording conditions (Casarotto *et al*., 2016). Although differences in TMS targets, stimulation intensity, EEG montages, and amplifier specifications across studies may introduce variability in absolute PCI values, the threshold has been validated across these variations, supporting its use as a consistent discriminative boundary. Although the PCI* = 0.31 threshold was originally derived using the PCI algorithm (Casarotto *et al*., 2016), it has been applied to PCIst values in subsequent studies (Comolatti *et al*., 2019). The present synthesis treats these algorithms as compatible for threshold-based classification, while acknowledging that absolute PCI values may differ slightly between methods.

Because the present synthesis is conceptual rather than meta-analytic, exact numerical pooling was not performed. Instead, each condition was positioned within the (M, C) space based on reported values relative to externally derived thresholds. This approach prioritises interpretive consistency over quantitative precision and is appropriate for evaluating the joint-regime prediction at a structural level. Nonetheless, differences in PET camera resolution, attenuation correction, region-of-interest definitions, TMS coil type, and preprocessing pipelines contribute to cross-study variance. Within-subject multimodal studies combining FDG-PET and TMS–EEG will be essential to validate the framework at the individual level.

## Results

### Joint metabolic–complexity structure of brain states

When mapped into the joint (M, C) space, canonical brain states resolve into a constrained distribution defined by both energetic availability and perturbational complexity (Figure 2; Tables 1–3). Across conditions, a consistent lower bound is evident along both axes. States associated with loss of consciousness fall below ∼42–46% of normal cortical metabolism (Stender *et al*., 2016) and exhibit reduced perturbational complexity (PCI < ∼0.31;(Casarotto *et al*., 2016)). In contrast, states associated with preserved or recovered awareness occupy a regime above both thresholds. This joint structure is consistent across independent cohorts despite differences in acquisition methods, analytical pipelines, and sampling frameworks. Notably, metabolism and perturbational complexity do not vary independently but co-occupy this restricted region across physiological, pharmacological, and clinical conditions.

**Figure 2.**
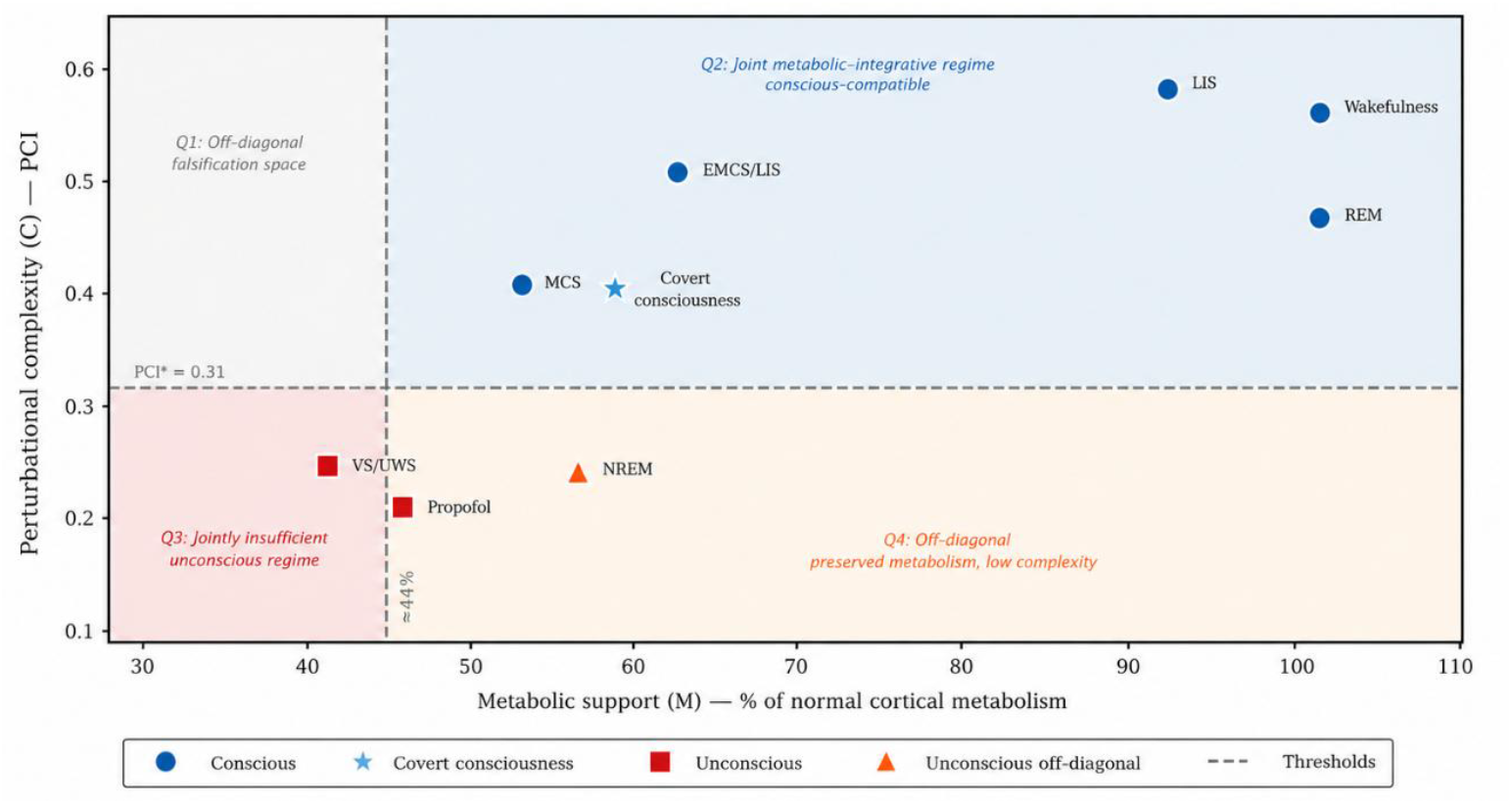
Distribution of canonical brain states. Fifteen canonical physiological, pharmacological, and clinical conditions mapped within the joint (M, C) state space. The metabolic axis (M) represents cortical glucose metabolism expressed as percentage of normal; the complexity axis (C) represents perturbational complexity (PCI). Dashed lines indicate the operational metabolic threshold (∼44% of normal cortical metabolism; lower bound of the empirically observed transition band ∼42–46%; Stender et al., 2016) and the complexity threshold (PCI* = 0.31; Casarotto et al., 2016), partitioning the space into four quadrants: Q2 (joint metabolic– integrative regime), Q3 (jointly insufficient regime), and Q1 and Q4 (off-diagonal, falsification-relevant regions). All 9 verified conscious states cluster within Q2, whereas all 6 unconscious states fall below at least one threshold (Q3 or Q4). The sole off-diagonal placement is NREM sleep (Q4), which exhibits preserved metabolic support (∼56% of normal) but reduced perturbational complexity (PCI ≈ 0.23), consistent with the prediction that metabolism alone is insufficient without preserved integrative dynamics. No verified conscious condition occupies an off-diagonal quadrant. Full values and sources are provided in Tables 1–3 and Supplementary Table S1.

### Metabolic axis

Across FDG-PET datasets, disorders of consciousness exhibit a graded distribution along the metabolic axis (Table 1;(Stender *et al*., 2016; Liu *et al*., 2024)). Vegetative state/unresponsive wakefulness syndrome (VS/UWS) consistently occupies a low-metabolic regime centred around ∼42% of normal cortical metabolism, whereas minimally conscious state (MCS) is associated with higher values (∼55–60%) (Stender *et al*., 2016). Transitional cases cluster near ∼44–46%, corresponding to the empirically observed boundary for recovery of behavioural responsiveness. Within the joint (M, C) space, these values position VS/UWS below the metabolic threshold associated with sustained large-scale integration, while MCS occupies a region compatible with partial restoration. Metabolic support alone, however, does not determine the organisation of neural activity, necessitating evaluation along the complexity axis (Table 1; Figure 2).

### Integrative (complexity) axis

Perturbational complexity measurements derived from TMS–EEG resolve conscious and unconscious states along a consistent boundary (Table 2) (Casali *et al*., 2013; Casarotto *et al*., 2016). Unconscious conditions—including deep sleep, anaesthesia, and VS/UWS—are associated with PCI values below ∼0.31 (Casarotto et al., 2016), reflecting reduced capacity for integrated yet differentiated cortical responses. In contrast, wakefulness, REM sleep, and minimally conscious states exhibit PCI values above this threshold (Casali *et al*., 2013; Casarotto *et al*., 2016), indicating preserved large-scale integrative dynamics. Within the joint (M, C) space, unconscious states fall below the integrative threshold, whereas conscious states occupy a regime characterised by sustained causal interactions across distributed networks. As with metabolism, however, complexity alone does not determine whether sufficient energetic support is available to maintain these dynamics (Table 2; Figure 2).

**Table 2.**
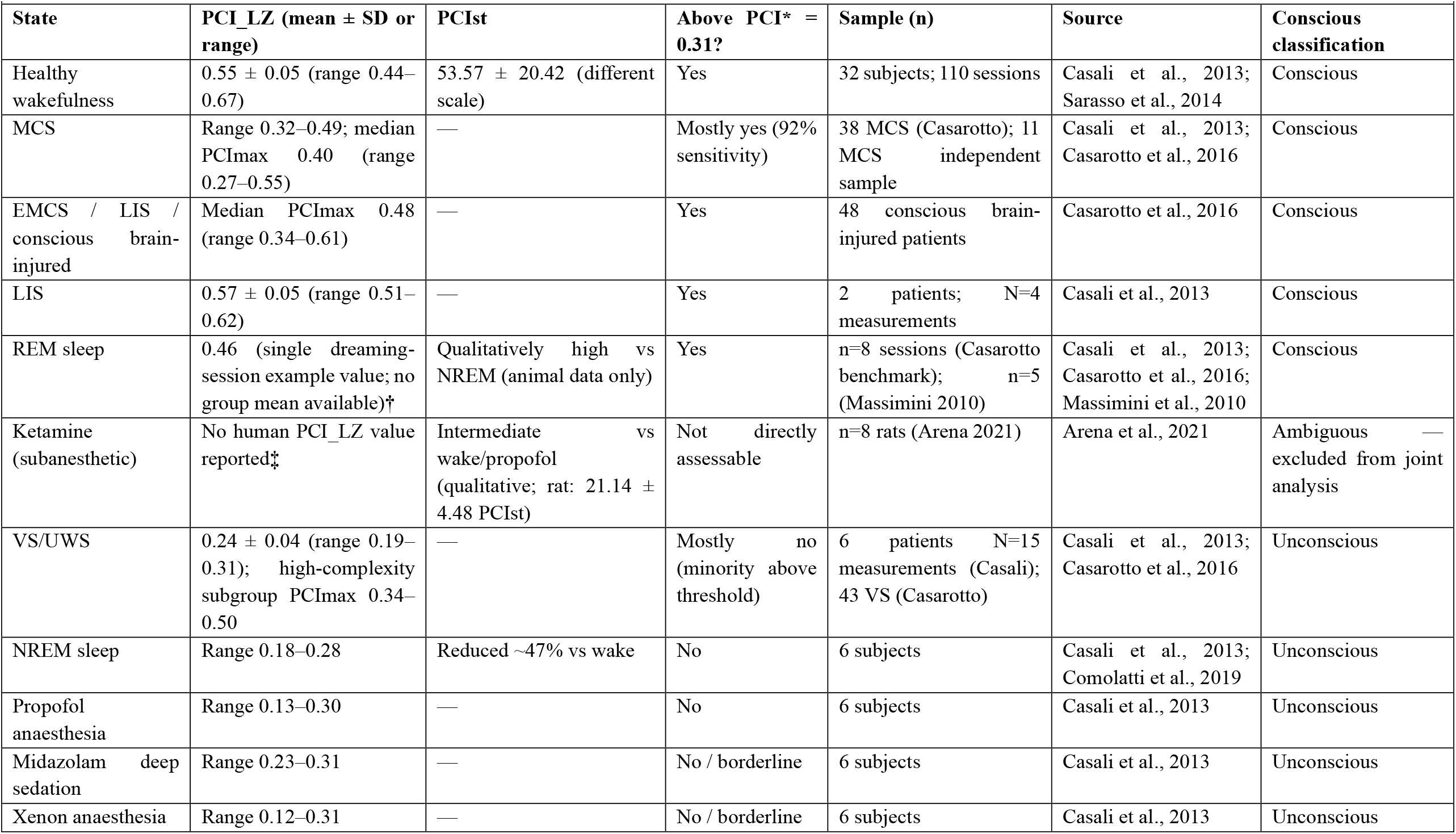
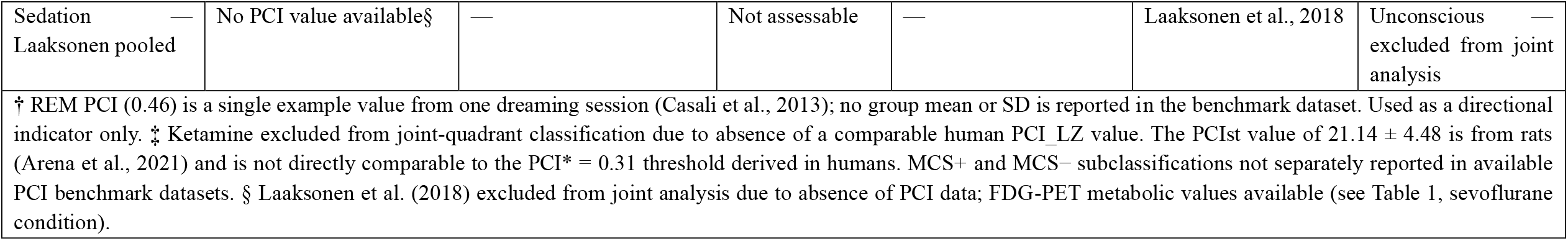
PCI / PCIst Values by Consciousness State.

| Table 2. PCI / PCIs Values by Consciousness State |  |  |  |  |  |  |
| --- | --- | --- | --- | --- | --- | --- |
| State | PCI_LZ (mean ± SD or range) | PCIs | Above PCI* = 0.31? | Sample (n) | Source | Conscious classification |
| Healthy wakefulness | 0.55 ± 0.05 (range 0.44–0.67) | 53.57 ± 20.42 (different scale) | Yes | 32 subjects; 110 sessions | Casali et al., 2013; Sarasso et al., 2014 | Conscious |
| MCS | Range 0.32–0.49; median PCImax 0.40 (range 0.27–0.55) | — | Mostly yes (92% sensitivity) | 38 MCS (Casarotto); 11 MCS independent sample | Casali et al., 2013; Casarotto et al., 2016 | Conscious |
| EMCS / LIS / conscious brain-injured | Median PCImax 0.48 (range 0.34–0.61) | — | Yes | 48 conscious brain-injured patients | Casarotto et al., 2016 | Conscious |
| LIS | 0.57 ± 0.05 (range 0.51–0.62) | — | Yes | 2 patients; N=4 measurements | Casali et al., 2013 | Conscious |
| REM sleep | 0.46 (single dreaming-session example value; no group mean available)† | Qualitatively high vs NREM (animal data only) | Yes | n=8 sessions (Casarotto benchmark); n=5 (Massimini 2010) | Casali et al., 2013; Casarotto et al., 2016; Massimini et al., 2010 | Conscious |
| Ketamine (subanesthetic) | No human PCI_LZ value reported‡ | Intermediate vs wake/propofol (qualitative; rat: 21.14 ± 4.48 PCIs) | Not directly assessable | n=8 rats (Arena 2021) | Arena et al., 2021 | Ambiguous — excluded from joint analysis |
| VS/UWS | 0.24 ± 0.04 (range 0.19–0.31); high-complexity subgroup PCImax 0.34–0.50 | — | Mostly no (minority above threshold) | 6 patients N=15 measurements (Casali); 43 VS (Casarotto) | Casali et al., 2013; Casarotto et al., 2016 | Unconscious |
| NREM sleep | Range 0.18–0.28 | Reduced ~47% vs wake | No | 6 subjects | Casali et al., 2013; Comolatti et al., 2019 | Unconscious |
| Propofol anaesthesia | Range 0.13–0.30 | — | No | 6 subjects | Casali et al., 2013 | Unconscious |
| Midazolam deep sedation | Range 0.23–0.31 | — | No / borderline | 6 subjects | Casali et al., 2013 | Unconscious |
| Xenon anaesthesia | Range 0.12–0.31 | — | No / borderline | 6 subjects | Casali et al., 2013 | Unconscious |
| Sedation —<br>Laaksonen pooled | No PCI value available§ | — | Not assessable | — | Laaksonen et al., 2018 | Unconscious —<br>excluded from joint analysis |
| † REM PCI (0.46) is a single example value from one dreaming session (Casali et al., 2013); no group mean or SD is reported in the benchmark dataset. Used as a directional indicator only. ‡ Ketamine excluded from joint-quadrant classification due to absence of a comparable human PCI_LZ value. The PCIst value of $21.14 \pm 4.48$ is from rats (Arena et al., 2021) and is not directly comparable to the $PCI^* = 0.31$ threshold derived in humans. MCS+ and MCS− subclassifications not separately reported in available PCI benchmark datasets. § Laaksonen et al. (2018) excluded from joint analysis due to absence of PCI data; FDG-PET metabolic values available (see Table 1, sevoflurane condition). |  |  |  |  |  |  |

### Joint regime

When metabolic and perturbational complexity measures are considered together, conscious states occupy a restricted region defined by the joint satisfaction of both constraints (Figure 2; Table 3). Wakefulness, REM sleep, and minimally conscious state (MCS) cluster within this regime, characterised by preserved metabolic support (>∼44–46%; Stender et al., 2016) and elevated perturbational complexity (PCI > ∼0.31; Casarotto et al., 2016). Within-cohort studies combining FDG-PET and TMS–EEG confirm this pattern, with MCS patients exhibiting both higher metabolism and PCI values above threshold relative to VS/UWS (Bodart *et al*., 2017), and multimodal FDG-PET/EEG assessments further improving diagnostic differentiation (Golkowski *et al*., 2017; Hermann *et al*., 2021).

**Table 3.** Paired Metabolic and Electrophysiological Markers Across States of Consciousness.

|  |  |  |  |  |  |  |
| --- | --- | --- | --- | --- | --- | --- |
| Study | Condition | Sample (n) | FDG-PET | PCI / EEG parameter | Key finding | Interpretation |
| Paired FDG-PET + PCI studies |  |  |  |  |  |  |
| Bodart et al., 2017 | VS/UWS | 9 | ~40–45% of normal | PCI < 0.31 | Low metabolism + low complexity | Unconscious regime (Q3) |
| Bodart et al., 2017 | MCS | 11 | ~55–60% of normal | PCI > 0.31 | High metabolism + high complexity | Joint regime — conscious-compatible (Q2) |
| Bodart et al., 2017 | EMCS / LIS (2 EMCS + 2 LIS) | 4 | Preserved (>60% of normal) | PCI > 0.31 | Both dimensions preserved | Conscious (Q2) |
| Bodart et al., 2017 | Behaviourally unresponsive subset† | 4 | Preserved (>55% of normal) | PCI > 0.31 | Preserved metabolism and complexity without behavioural output | Within joint regime — covert consciousness (Q2) |
| Paired FDG-PET + other EEG markers |  |  |  |  |  |  |
| Hermann et al., 2021 | VS/UWS vs MCS | 52 (21 VS + 31 MCS) | MIBH (not normalised to % of normal cortical metabolism); AUC 0.821 for MCS diagnosis | EEG SVM classifier on 112 spectral / connectivity / complexity / ERP features | PET-MIBH + EEG combination: 94% sensitivity, 67% specificity for MCS | Multimodal differentiation outperforms either modality alone |
| Hermann et al., 2021 | VS/UWS with covert neural correlate of consciousness | 4 of 21 VS patients | Preserved MIBH in covert subset | Concordant EEG-positive on local-global paradigm | 4/7 (57%) of VS patients with complete multimodal data showed neural correlates of consciousness vs 1/14 (7%) without (p = 0.025); 9/24 (38%) of initially unresponsive patients recovered command-following at 6 months vs 0/16 (0%, p = 0.006) | Candidate covert consciousness (Q2 placement consistent with the joint-regime framework) |

| Study | Condition | Sample (n) | FDG-PET | PCI / EEG parameter | Key finding | Interpretation |
| --- | --- | --- | --- | --- | --- | --- |
| Golkowski et al., 2017 | VS/UWS vs MCS | 20 | Occipital metabolism higher in MCS than VS/UWS | Delta band power | PET–EEG concordance differentiates states | Reduced integration in VS/UWS |

FDG-PET: fluorodeoxyglucose positron emission tomography; PCI: perturbational complexity index; VS/UWS: vegetative state/unresponsive wakefulness syndrome; MCS: minimally conscious state; EMCS: emergence from minimally conscious state; LIS: locked-in syndrome; MIBH: metabolic index of the best-preserved hemisphere; SVM: support vector machine; ERP: event-related potential.
Reference boundaries: metabolic support ~42–46% of normal cortical metabolism (operational cutoff ~44%); PCI\* ≈ 0.31. The upper section presents studies with paired FDG-PET and PCI measurements; the lower section presents studies combining FDG-PET with other electrophysiological markers. Only PCI-based measurements are directly comparable to the PCI\* threshold used in the joint-regime analysis. EEG-based markers in the lower section inform multimodal diagnostic accuracy but are not used for quadrant classification. † Cases with preserved metabolism and PCI above PCI\* in behaviourally unresponsive patients do not violate the joint metabolic–integrative constraint; they indicate covertly preserved conscious capacity not accessible through behavioural output.

In contrast, unconscious states—including deep anaesthesia and vegetative state/unresponsive wakefulness syndrome (VS/UWS)—fall outside this region along one or both axes. Propofol-induced loss of consciousness is associated with coordinated reductions in metabolism and complexity, producing trajectories that cross the joint boundary (Alkire *et al*., 1995; Laaksonen *et al*., 2018). Disorders of consciousness similarly distribute below one or both thresholds, depending on clinical state.

The empirical distribution is further defined by the fact that no verified conscious state occupies an off-diagonal position in currently available datasets, although the limited availability of paired within-subject measurements means this should be interpreted as a current empirical observation rather than definitive exclusion. States combining high perturbational complexity with insufficient metabolic support, or preserved metabolism with reduced complexity, are under-represented, with the only off-diagonal placement corresponding to NREM sleep — an unconscious state (Figure 2). These off-diagonal regions constitute critical empirical tests of the joint-regime prediction.

### Informative state transitions

Transitions between conscious and unconscious states correspond to boundary crossings within the joint (M, C) space rather than to variation along a single axis. The transition from VS/UWS to minimally conscious state (MCS) is associated with concurrent increases in both metabolism and perturbational complexity, shifting the system into the joint regime. Conversely, anaesthetic induction produces coordinated reductions along both axes, resulting in exit from this regime.

REM sleep provides a critical test case. Despite behavioural disconnection from the external environment, it exhibits high perturbational complexity (Casali *et al*., 2013). Although paired metabolic measurements are limited, available evidence indicates that global metabolism remains above levels associated with unconscious states, placing REM within or near the joint regime. This dissociation demonstrates that environmental responsiveness is not a necessary condition for preserved integrative capacity. Direct within-subject measurements would determine whether REM occupies a distinct subregion within the joint space.

Together, these transitions support the interpretation that conscious states are defined by a constrained region of jointly sufficient metabolic and integrative conditions, rather than by either variable in isolation.

### Empirical constraints and falsification space

When mapped into the joint (M, C) state space, the predicted “joint-regime” region is defined by the conjunction of (i) sufficient metabolic support (M ≥ ∼44% of normal cortical metabolism) and (ii) preserved perturbational complexity (C ≥ PCI* = 0.31). Accordingly, four quadrants can be distinguished: Q1 (M < threshold, C ≥ PCI*; low metabolism, high complexity; falsification space), Q2 (M ≥ threshold, C ≥ PCI*; the joint metabolic–integrative regime), Q3 (M < threshold, C < PCI*; jointly insufficient regime), and Q4 (M ≥ threshold, C < PCI*; preserved metabolism but reduced complexity; falsification space).

Across the 18 canonical conditions initially mapped (Tables 1–3; Figure 2), 15 had complete data on both axes and were included in the joint-regime analysis. Two conditions — midazolam anaesthesia and xenon anaesthesia — were excluded because no quantitative FDG-PET metabolic values were identified in the available literature for these agents (see Methods; Supplementary Table S1). A further sedation condition from Laaksonen et al. (2018) was excluded because corresponding PCI measurements were unavailable. Of the 15 conditions with complete (M, C) data, 9 occupied Q2, 1 occupied Q4, 0 occupied Q1, and 5 occupied Q3.

One condition — propofol anaesthesia — sits near the metabolic threshold (M ≈ 45%, marginally above the 44% operational cutoff; C ≈ 0.21, below PCI*). Because the propofol metabolic value is derived from an absolute CMRGlu ratio (Alkire et al., 1995) rather than the same percentage-of-control normalisation used for disorders-of-consciousness benchmarks, it was conservatively classified as Q3 in the primary analysis. A sensitivity analysis reassigning propofol to Q4 did not alter the overall pattern: all 9 conscious conditions remained within Q2, no conscious condition occupied an off-diagonal quadrant, and Fisher’s exact test remained significant (two-tailed p < 0.001). The conclusion is therefore robust to this boundary assignment (see Supplementary Table S1, footnote on propofol classification).

NREM sleep (N2–N3) occupies Q4: its metabolism (∼56% of normal;(Maquet *et al*., 1990)) exceeds the metabolic threshold, but its perturbational complexity (∼0.23; (Casali *et al*., 2013)) falls below PCI*. This placement does not constitute a falsifying observation because NREM sleep is an unconscious state. The framework predicts that conscious experience should not occur in off-diagonal quadrants; it does not predict that all off-diagonal configurations are impossible, only that they should not be accompanied by verified awareness. Indeed, NREM sleep in Q4 provides positive evidence for the framework: it demonstrates that metabolic support alone is insufficient to sustain conscious experience in the absence of preserved perturbational complexity.

Importantly, both analyses treat condition-level, cross-study estimates as observational units rather than independent individual-level data and should therefore be interpreted as descriptive summaries of distributional structure rather than conventional inferential tests. Descriptively, the 2 × 2 classification (consciousness status versus joint threshold satisfaction) yielded a perfectly diagonal pattern: all 9 conscious conditions met both thresholds, whereas none of the 6 unconscious conditions met both (Fisher’s exact test, two-tailed p = 0.0002). In parallel, metabolic support and perturbational complexity exhibited a strong positive monotonic association across the 15 conditions with complete (M, C) data (Spearman’s ρ = 0.95, p < 0.001).

These findings yield a clear falsification target for future work. Verified conscious experience in either Q1 (low metabolism with preserved complexity) or Q4 (preserved metabolism with reduced complexity) would directly challenge the joint-regime framework. The present mapping is derived from cross-study, group-level estimates, and within-subject datasets combining quantitative FDG-PET metabolism and TMS–EEG remain limited. Consequently, the absence of conscious states in off-diagonal regions may reflect either a genuine biological constraint or a limitation of current measurement coverage. Prospective multimodal studies integrating FDG-PET and TMS–EEG within individual subjects will be critical for determining whether the joint-regime structure is preserved at the individual level and for populating the currently under-sampled off-diagonal regions.

## Discussion

The present analysis identifies a structured relationship between metabolic support and perturbational complexity that constrains the conditions under which conscious states are observed. When represented within a joint (M, C) space, canonical brain states do not distribute freely but cluster within a bounded region defined by sufficient energetic availability and preserved large-scale integrative dynamics. This structure is consistently observed across physiological, pharmacological, and clinical conditions.

This joint organisation reframes prior findings. Measures of perturbational complexity reliably distinguish conscious from unconscious conditions across wakefulness, sleep, anaesthesia, and disorders of consciousness, yet do not specify the energetic requirements necessary to sustain such dynamics. Conversely, FDG-PET studies identify metabolic thresholds associated with recovery of awareness (Stender *et al*., 2016) but do not resolve the organisational properties of neural activity within those regimes. Considered together, these lines of evidence define complementary constraints that jointly delimit the space of viable conscious states.

The observed structure is best explained by a joint metabolic–integrative constraint: conscious states arise only when both sufficient metabolic support and preserved perturbational complexity are simultaneously satisfied. This formulation imposes a falsifiable exclusion: reportable experience should not occur outside this regime. In this view, consciousness is not attributed to a specific mechanism or region, but to a regime of system-level organisation that becomes available only under defined energetic and dynamical conditions.

Within this framework, transitions between states are most naturally interpreted as boundary crossings in a constrained state space rather than as variation along a single axis. The transition from VS/UWS to minimally conscious state (MCS) is associated with concurrent increases in both metabolism and perturbational complexity, while anaesthetic induction produces coordinated reductions along both axes. These transitions reflect entry into and exit from a jointly sufficient regime, providing a unified account of otherwise heterogeneous phenomena across disorders of consciousness, pharmacological manipulation, and physiological state changes.

A central empirical feature of the present mapping is that no verified conscious state has yet been observed to occupy an off-diagonal position — the only such placement, NREM sleep in Q4, is unconscious. This observation has two implications. First, it indicates that the two axes, while strongly co-varying, are not redundant: metabolic support can be preserved while integrative complexity collapses, yet consciousness does not follow metabolism alone. Second, it sharpens the falsification criterion to a specific empirical target: verified conscious experience in Q1 or Q4 would directly challenge the joint-regime prediction. Whether the current absence of such cases reflects a genuine biological constraint or incomplete sampling of the state space remains the key open question.

The strong positive association between metabolic support and perturbational complexity across conditions (Spearman’s ρ = 0.95) reflects not redundancy but a shared enabling structure: both constraints must be jointly satisfied, and their co-variation is a consequence of this rather than evidence that one reduces to the other.

At present, however, this mapping is derived primarily from cross-study and group-level estimates, as within-subject datasets combining quantitative FDG-PET metabolism and perturbational complexity remain limited (but see (Bodart *et al*., 2017; Hermann *et al*., 2021)). Consequently, the absence of conscious states in off-diagonal configurations may reflect either a genuine biological constraint or a limitation of current measurement coverage. Direct within-subject multimodal studies integrating FDG-PET and TMS–EEG will be essential to resolve this ambiguity.

Clinical dissociation is a further motivation for multimodal neuroimaging: impaired behavioural responsiveness need not map cleanly onto the presence or absence of residual brain function, because patients can have sensory-motor deficits that differentially limit behavioural capacity and thereby complicate bedside inference about conscious state. This clinical dissociation motivates the use of neuroimaging to identify residual brain functions when overt behaviour is compromised, and it provides a practical rationale for frameworks that treat consciousness as constrained by measurable biological conditions rather than by behavioural output alone.

REM sleep represents an informative intermediate case. It exhibits high perturbational complexity despite behavioural disconnection from the external environment (Massimini *et al*., 2005; Casali *et al*., 2013). Critically, FDG-PET studies indicate that cerebral glucose metabolism during REM is comparable to wakefulness: Maquet et al. (1990) reported that global cortical metabolic rate during REM did not differ significantly from waking values (mean difference: +4.30 ± 7.40%, P = 0.35), whereas slow-wave sleep showed a substantial reduction (−43.80 ± 14.10%, P < 0.01). Consistent with preserved global energetic support, preferential activation of limbic and paralimbic structures during REM sleep has been demonstrated using regional cerebral blood flow imaging, with perfusion in these regions exceeding waking levels (Braun *et al*., 1997). Although regional cerebral blood flow is not a direct measure of glucose metabolism, it is commonly interpreted as a proxy under conditions of preserved neurovascular coupling. Elevated whole-brain metabolism during REM relative to wakefulness has also been observed, particularly in centrencephalic and limbic regions (Nofzinger *et al*., 1997). Together, these metabolic findings place REM well above the ∼42– 46% metabolic threshold associated with unconscious states (Stender *et al*., 2016), supporting its placement within or near the joint regime. However, definitive placement requires paired within-subject measurement of both PCI and FDG-PET, making REM a critical test condition for the joint-regime framework.

Ketamine provides a complementary test case. Subanesthetic ketamine induces dissociative conscious experiences while altering both metabolic profiles (Långsjö *et al*., 2004) and TMS-evoked dynamics (Sarasso *et al*., 2015). Such states may occupy boundary or transitional regions within the (M, C) space. Precise placement of ketamine conditions therefore represents a priority for future within-subject studies aimed at probing the limits of the joint constraint.

The present framework is complementary to existing theories of consciousness but differs in its emphasis on the biological constraints that enable the computations each theory describes. Integrated Information Theory (Tononi, 2004) formalises consciousness in terms of integrated information (Φ), yet the causal architecture over which Φ is computed depends on sustained synaptic interactions that require continuous metabolic support; the present framework makes this energetic dependency explicit. Global Workspace Theory (Baars, 1988; Dehaene and Changeux, 2011) characterises consciousness as large-scale broadcasting of selected information across distributed cortical networks, but such widespread cortical activation is itself energetically demanding, and the metabolic axis operationalises this requirement by identifying the lower bound of energetic availability below which global broadcasting cannot be sustained. Higher-order theories (Lau and Rosenthal, 2011) emphasise that conscious perception requires higher-order representational states, which in turn depend on functional prefrontal circuits that are among the most metabolically demanding regions of the cortex. Rather than competing with these accounts, the current approach identifies shared enabling conditions: the metabolic and integrative constraints within which the mechanisms posited by each theory can operate. In this sense, the joint-regime framework specifies the biological regime that must be satisfied for integrated information, global broadcasting, or higher-order representation to be physically instantiated.

An important alternative account is that the observed co-variation between metabolism and perturbational complexity reflects shared dependence on global arousal rather than a genuine joint biological constraint. Because arousal modulates both cortical metabolic rate and the capacity for sustained causal interactions, the constrained distribution within the (M, C) space could in principle arise as an epiphenomenon of arousal-level variation. However, several observations argue against a purely arousal-driven explanation. First, REM sleep exhibits high perturbational complexity (Casali *et al*., 2013) and preserved global metabolism (Maquet *et al*., 1990) despite markedly reduced behavioural arousal, demonstrating that the joint regime can be occupied independently of arousal state. Conversely, patients in unresponsive wakefulness syndrome (VS/UWS) exhibit preserved sleep–wake cycling—and thus intact arousal mechanisms—yet fall below both metabolic and complexity thresholds, indicating that arousal alone is insufficient to place a state within the joint regime. Beyond arousal, the disorders-of-consciousness literature introduces additional confounds including medication effects, aetiology, injury severity, and time post-injury, all of which can independently affect both FDG-PET and TMS–EEG measurements at the group level. The joint-regime framework generates a specific prediction regarding these confounds: if the metabolic–integrative constraint is genuine, then inter-individual variation in medication status or injury characteristics should not systematically produce off-diagonal configurations—that is, states with high complexity under insufficient metabolism, or preserved metabolism with reduced complexity. Resolving this prediction will require within-subject multimodal designs in which FDG-PET and TMS–EEG are acquired in the same individuals, thereby controlling for the compositional differences inherent in cross-study synthesis.

Finally, available metabolic evidence suggests that consciousness may be constrained by an energetic boundary: the finding that ∼42% of normal cortical activity can constitute a minimal energetic requirement for conscious awareness suggests that below a certain level of global metabolic support, conscious awareness is unlikely to be sustained. This kind of boundary result provides a biologically grounded constraint on which brain states can plausibly support conscious experience, especially in clinical contexts where behaviour is an unreliable proxy for awareness.

More broadly, these findings suggest that persistent lack of convergence in consciousness research (Seth and Bayne, 2022) may reflect a mismatch between the level at which explanations are formulated and the level at which biological constraints operate. By framing consciousness as a regime within a constrained state space, the problem shifts from identifying a singular mechanism to characterising the conditions under which particular dynamical organisations become possible.

## Conclusion

Conscious states occupy a constrained region of state space defined by the joint satisfaction of sufficient metabolic support and preserved perturbational complexity. This structure is consistently observed across physiological, pharmacological, and clinical conditions and is not captured by either dimension in isolation. The absence of verified conscious experience in off-diagonal configurations — high complexity without sufficient metabolic support or preserved metabolism with reduced complexity — defines a falsifiable prediction: reportable experience should not occur outside the joint metabolic–integrative regime. Whether this absence reflects a fundamental biological constraint or a limitation of current measurement remains an open question. Direct within-subject multimodal investigations integrating FDG-PET and TMS– EEG across controlled state transitions will be critical in determining whether the observed structure represents a general principle of brain organisation. In this framework, consciousness is not a property of a specific mechanism, but a regime of biological organisation — and identifying its boundaries offers both a principled target for future neuroscience and a practical basis for assessing residual awareness in patients who cannot report their experience behaviourally.

## Author Contributions

T.S.: Conceptualization, Methodology, Investigation, Formal analysis, Writing — original draft, Visualization. A.M.: Writing — review and editing. Both authors read and approved the final manuscript.

## Data Availability Statement

This study is a constraint-driven cross-study synthesis; no new primary data were generated. All quantitative values used in the joint metabolic–integrative analysis were extracted from previously published, peer-reviewed sources, which are individually cited in the main text and listed with their original references in Tables 1–3 and Supplementary Table S1. The complete cross-study dataset used for the joint-regime classification — including all (M, C) coordinates, per-condition source attributions, and quadrant assignments — is provided in Supplementary Table S1 and is additionally archived in the associated Open Science Framework project at [OSF DOI — to be inserted on deposit]. No analysis code beyond the descriptive statistics reported in the main text (Fisher’s exact test, Spearman’s rank correlation) was required; figure-generation scripts are available from the corresponding author on reasonable request.

## Related Work

This manuscript is one component of a coordinated research programme — the Constraint-Based Bayesian Residue Approach (COBRA) — which develops a multiscale, constraint-based framework spanning consciousness, mental health, systems-based medicine, and artificial systems. Companion preprints addressing related components of the programme are listed in the reference section. Each paper is intended to stand alone as a falsifiable contribution; cross-references are provided where one paper’s claims depend on another’s.

## Ethics Statement

This study did not involve human participants, human data, or animal subjects. Empirical findings discussed in this manuscript are drawn from previously published sources, each of which carries its own ethics approval as reported in the original study.

## AI Assistance Statement

AI assistance was used during manuscript preparation, using Claude (Anthropic). Specifically, the tool was used for language editing, reference verification, drafting of standard back-matter sections (Data Availability, AI Assistance, related-work disclosures), and assistance with figure codes. No AI tool authored scientific content, generated novel claims, designed the framework, analysed data, or made any interpretive judgements. All scientific content, interpretations, and conclusions are the work of the authors, who take full responsibility for the manuscript.

---

**Supplementary Table S1.** Complete (M, C) dataset for the joint-regime analysis.

|  |  |  |  |  |  |  |  |  |  |
| --- | --- | --- | --- | --- | --- | --- | --- | --- | --- |
| # | Condition | Conscious | M (% normal) | M source | M notes | C (PCI) | C source | C notes | Quadrant |
| 1 | Healthy wakefulness | Yes | 100% | Stender et al., 2016 | Defined as normative baseline; all DoC values normalised to this | 0.55 (range 0.44–0.67) | Casali et al., 2013 | 32 awake healthy subjects; 110 sessions | Q2 |
| 2 | REM sleep | Yes | ~100% | Maquet et al., 1990 | Global metabolic rate not significantly different from waking (+4.30 ± 7.40%, p = 0.35) | 0.46 (single dreaming-session example value) | Casali et al., 2013 | No group mean available; single session within conscious distribution | Q2 |
| 3 | MCS (Stender / Casarotto) | Yes | 55% | Stender et al., 2016 | Global cortical CMRglc 55% of normal; n = 21 MCS | 0.40 (median PCI<sub>max</sub>; range 0.32–0.49) | Casarotto et al., 2016 | PCI<sub>max</sub> > PCI* in 92% of MCS; n = 38 | Q2 |
| 4 | EMCS (Stender / Casarotto) | Yes | ~55% | Stender et al., 2016 | Metabolic rates indistinguishable from MCS in source; no separate % of normal reported | 0.48 (median PCI<sub>max</sub>; range 0.34–0.61) | Casarotto et al., 2016 | 48 conscious brain-injured patients including EMCS, LIS, stroke | Q2 |
| 5 | LIS (Casali) | Yes | ~90% (near-normal) | Soddu et al., 2015 | Gray-matter SUV 5.6 ± 3.2 vs controls 6.0 ± 0.9; not expressed as % of normal; assumed near-normal | 0.57 (range 0.51–0.62) | Casali et al., 2013 | 2 LIS patients; N = 4 measurements | Q2 |
| 6 | MCS (Bodart 2017) | Yes | ~57% (~55–60%) | Bodart et al., 2017 | Directional; from paired FDG-PET + TMS–EEG study | 0.40 (> PCI* = 0.31) | Bodart et al., 2017 | Same paired study; PCI<sub>max</sub> > 0.31 confirmed | Q2 |
| 7 | EMCS / LIS (Bodart 2017) | Yes | ~62% (>60%) | Bodart et al., 2017 | Preserved metabolic rates; includes 2 LIS + 2 EMCS patients | 0.50 (> PCI* = 0.31) | Bodart et al., 2017 | High complexity confirmed; same paired study | Q2 |
| 8 | Covert consciousness subset (Bodart 2017) | Yes (covert) | ~57% (>55%) | Bodart et al., 2017 | 4 behaviourally unresponsive patients with preserved metabolic rates | 0.40 (> PCI* = 0.31) | Bodart et al., 2017 | High complexity confirmed despite behavioural unresponsiveness | Q2 |
| 9 | MCS (Liu 2024) | Yes | 53% | Liu et al., 2024 | Mean cortical metabolic index 4.19 vs controls 7.88 = 53%; n = 87 MCS | 0.40 (cross-mapped from Casarotto 2016) | Casarotto et al., 2016 | C value cross-mapped from matched diagnostic category; not paired in Liu 2024 | Q2 |
| 10 | VS/UWS (Stender / Casali) | No | 42% | Stender et al., 2016 | Global cortical CMRglc 42% of normal; n = 14 VS/UWS | 0.24 (mean ± 0.04; range 0.19–0.31) | Casali et al., 2013 | 6 VS/UWS patients; N = 15 measurements | Q3 |
| 11 | VS/UWS (Casarotto 2016) | No | 42% | Stender et al., 2016 (cross-mapped) | M not independently measured in Casarotto 2016; cross-mapped from Stender diagnostic class | 0.25 (median PCI<sub>max</sub>; range 0.19–0.31) | Casarotto et al., 2016 | 43 VS patients; 9 high-complexity subgroup (PCI<sub>max</sub> 0.34–0.50) not counted here | Q3 |
| 12 | VS/UWS (Liu 2024) | No | 41% | Liu et al., 2024 | Best-fit threshold 3.32 = 41% of healthy mean 7.88; n = 48 VS/UWS | 0.24 (cross-mapped from Casali 2013) | Casali et al., 2013 | C cross-mapped from matched diagnostic category; not paired | Q3 |
| 13 | VS/UWS (Bodart 2017) | No | ~42% (~40–45%) | Bodart et al., 2017 | Directional; low metabolism confirmed in paired study | 0.25 (< PCI* = 0.31) | Bodart et al., 2017 | Low complexity confirmed; congruent with metabolic classification | Q3 |
| 14 | Propofol anaesthesia | No | ~45% | Alkire et al., 1995 | Awake GMR 29 ± 8; anaesthetised 13 ± 4 $\mu\text{mol}\cdot 100\text{g}^{-1}\cdot \text{min}^{-1}$; ratio = 45%; not normalised to separate control group | 0.21 (midpoint; range 0.13–0.30) | Casali et al., 2013 | 6 healthy volunteers; propofol titrated to unresponsiveness | Q3* |
| 15 | NREM sleep | No | ~56% | Maquet et al., 1990 | Global rate reduced by $-43.80 \pm 14.10\%$ vs waking ($p < 0.01$), yielding ~56% of waking level | 0.23 (midpoint; range 0.18–0.28) | Casali et al., 2013 | 6 healthy subjects progressing from wakefulness to NREM | Q4 |

\* Propofol M = 45% falls marginally above the 44% operational threshold, whereas C = 0.21 falls below PCI\*. Under strict threshold application, this would place propofol in Q4. However, because the metabolic value is derived from an absolute CMRglu ratio rather than the same normalisation framework used for disorders-of-consciousness conditions, it was conservatively classified as Q3 in the primary analysis. Sensitivity analyses treating propofol as Q4 did not alter the overall pattern: all 9 conscious conditions remained within Q2, and Fisher's exact test remained significant ( $p < 0.001$ ). † NREM sleep is the only off-diagonal condition (M above threshold; C below PCI\*) and is unconscious, consistent with the joint-constraint prediction that metabolic support alone is insufficient for consciousness in the absence of preserved perturbational complexity.

